# Speeding up, not slowing down, decreases the energy needed to control walking balance

**DOI:** 10.1101/2025.03.25.645241

**Authors:** Wouter Muijres, Maarten Afschrift, Renaud Ronsse, Friedl De Groote

## Abstract

There is a metabolic cost associated with controlling walking balance but it remains unclear to what extent sagittal plane balance control contributes to this cost. Furthermore, sagittal plane balance control strategies vary with speed but it is unclear whether this also leads to a speed-dependent balance-related metabolic cost. Here, we explored the metabolic cost of stabilizing walking in the sagittal plane across speeds and its relationship with balance control strategies. To this aim, we applied continuous treadmill belt speed perturbations (standard deviation of 0.13 ms^−1^) to 22 healthy individuals walking at 0.8, 1.2, and 1.6 m/s. We evaluated changes in metabolic energy consumption and balance control strategies between perturbed and unperturbed walking and explored relationships between both. Perturbations induced larger increases in metabolic rate and changes in balance control strategies at slower than faster walking speeds, suggesting that walking is more robust against perturbations at faster speeds. Perturbations increased the metabolic rate by 16.7% at the slowest versus 4.6% at the fastest walking speed. When perturbed, subjects took shorter, wider, and more variable steps, and variability in ankle muscle activation increased but most changes were larger at slower speeds. Metabolic rate increased more due to perturbations in individuals who reduced step length more, i.e. relied more on anticipatory adjustments of the walking pattern. Our findings are especially relevant for explaining the increased metabolic cost of individuals with mobility impairments, who often walk slower and have altered balance control.

## Introduction

There is a metabolic cost associated with controlling walking balance. The relationship between metabolic cost and balance control has been studied in the frontal plane by either stabilizing balance or perturbing sensory input using visual perturbations (1–3). Increases in the metabolic cost of walking were associated with both anticipatory adjustments in step length or step width (3) and reactive adjustments in foot placement resulting in increased step length and step width variability (2). However, for stabilizing walking in the sagittal plane, healthy adults also rely on reactive ankle torques, i.e. the ankle strategy (4). The ankle strategy is used more at slow than at fast speeds (4). Yet, it is unclear whether individuals rely less on the ankle strategy at fast walking speeds because they rely on other strategies or because they require less active balance control to maintain stability because walking is more robust against perturbations at higher speeds (5, 6). In the first case, the metabolic cost for stabilizing walking at fast speeds would not necessarily be lower if anticipatory or reactive foot placement strategies are indeed important determinants of metabolic cost, whereas in the second case, we would expect a lower cost for stabilizing walking at fast speeds. Here, we explored the metabolic cost of stabilizing walking in the sagittal plane across speeds and its relationship with balance control strategies.

### The metabolic cost of controlling walking balance has been studied experimentally by either increasing or reducing the need for balance control

When reducing the need for frontal plane balance control by springs that stabilize the pelvis in the mediolateral direction, energy consumption decreased by 5.7% compared to walking without stabilization (1). On the other hand, mediolateral virtual perturbations to the visual field increased walking energy consumption by 5.9%-12% compared to unperturbed walking (2, 3). Most studies focused on frontal plane balance control because walking is thought to be more unstable in the frontal plane (1). In addition, the cost of walking on uneven terrain, which challenges balance control across planes, has been studied. When walking on uneven surfaces, healthy adults increased energy consumption by 28% (7). Together, these studies suggest that there is a considerable energetic cost associated with walking balance control, but it remains unclear if, and to what extent, sagittal plane balance control contributes to this cost.

### Humans use both anticipatory and reactive balance control strategies

Healthy adults increase step width and decrease step length in the presence of continuous perturbations (2, 3, 7, 8) or when expecting discrete perturbations whose timing is unpredictable (9, 10). In addition, humans use reactive stepping and ankle strategies. Healthy adults adapt foot placement in response to disturbances such as uneven terrain (e.g. 7), mechanical (e.g. 11, 13), and visual perturbations (2, 3, 8). Healthy adults also restore balance by modulating their ankle torque by adapting ankle muscle activity to control the center of pressure. This ankle strategy is mainly used in the sagittal plane where the shape of the foot allows for larger displacements of the center of pressure (4, 13). In summary, healthy adults stabilize walking against disturbances by combining anticipatory adjustments including increases in step width and reductions in step length, with reactive foot placement and ankle responses.

### The energetic cost of walking balance has been related to both anticipatory adjustments in mean step length and step width, and reactive foot placement strategies

When stabilizing the pelvis in the frontal plane, Donelan et al. (1) observed both a decrease in energetic cost and in step width variability and, therefore, suggested that the reduced need for using a reactive foot placement strategy in the frontal plane was the most likely explanation for the observed reductions in energy cost. O’Connor et al. (2) tested associations between increases in energy consumption due to visual field perturbation and changes in anticipatory (mean step width) and reactive (step width variability) stepping parameters. They found that energetic cost was most strongly associated with step width variability. However, Ahuja and Franz (3) observed simultaneous changes in energetic cost and anticipatory adjustments in step length and width. Healthy adults initially took shorter and wider steps and consumed more energy in response to mediolateral perturbations of the visual field, but prolonged exposure to perturbations resulted in less anticipatory adjustments and a 5% decrease in energy consumption. Hence, both alterations in anticipatory and reactive stepping strategies might have an important contribution to the energetic cost of frontal plane balance control. It is unclear whether this is also true for sagittal plane balance control where the reliance on the ankle strategy is larger.

### Healthy adults rely more on an ankle strategy to control sagittal plane balance at slower than faster speeds

Modulation of ankle moments in response to treadmill belt speed perturbations is larger during slow compared to fast walking (4). It is unclear whether a smaller reliance on the ankle strategy at faster speeds is associated with a larger reliance on stepping strategies. In the frontal plane, mediolateral foot placement in unperturbed walking is better explained by center of mass kinematics at faster walking speeds, suggesting an increased reliance on foot placement strategies (6). It is possible that something similar happens in the sagittal plane. With increasing speed, step frequency increases, and thus, the time between consecutive foot contacts decreases, allowing for faster corrections through foot placement (4). At the same time, reductions in stance time decrease the available time for moving the center of pressure under the foot and, thus, the application of an ankle strategy. However, it has also been suggested that passive walking dynamics is more robust against perturbations in the sagittal plane at faster speeds (5, 6). Whether sagittal plane balance control relies more on stepping or is easier at faster speeds will have consequences for how the energetic cost of sagittal plane balance control varies with speed.

### In this study, we assessed the metabolic cost of stabilizing walking in the sagittal plane across speeds and explored its relationship with balance control strategies

To this aim, we applied continuous treadmill belt speed perturbations to healthy individuals. Such perturbations challenge walking balance control in the sagittal plane and were previously found to elicit ankle responses, which were stronger at slower speeds (4). We hypothesized that these perturbations would increase the energy consumption of walking balance control and that increases in energy consumption would be speed-dependent. In addition, we explored whether between-subject variation in energy consumption for balance control could be explained by variation in anticipatory and reactive balance control strategies. We assessed anticipatory alterations in the gait pattern based on changes in average step length and step width. We assessed reactive foot placement strategies based on changes in step length and step width variability, and reactive ankle strategies based on changes in variability of ankle muscle activity (measured through electromyography).

## Methods

### Description subjects

We recruited subjects who understood English or Dutch. Individuals were excluded from participation when they suffered from conditions or took medication that influenced balance control or locomotor ability. We collected a representative dataset by including at least one female and one male participant in each age range (18-20, 21-25, 26-30, …, 56-60, and 61-65 years old). In total, 22 healthy adults between 19-65 years old (female: N=12, mean body mass: 71.9 kg, mean leg length: 0.93 m) enrolled in this study and provided written informed consent. The experimental protocol was approved by the Social and Societal Ethics Committee of the KU Leuven (G-2024-7626). The experimental protocol was pre-registered on Open Science Framework (14) after collecting data from the first three subjects.

### Experimental procedure

Subjects walked on an instrumented treadmill (M-Gait, Motekforce link, Amsterdam, Netherlands) at three speeds, 0.8, 1.2, and 1.6 ms^−1^, with and without treadmill belt speed perturbations. Walking conditions were randomized per block, i.e. first by speed and then by perturbation condition (Figure 1A). In each trial, subjects walked for about 6 minutes with at least 5 minutes of rest between walking trials. Subjects performed a 5-minute unperturbed walking trial for habituation at 1.2 ms^−1^ before the start of the experiment. During walking trials, subjects wore a safety harness, tethered to the ceiling, to prevent falls in the event of balance loss.

**Figure 1.**
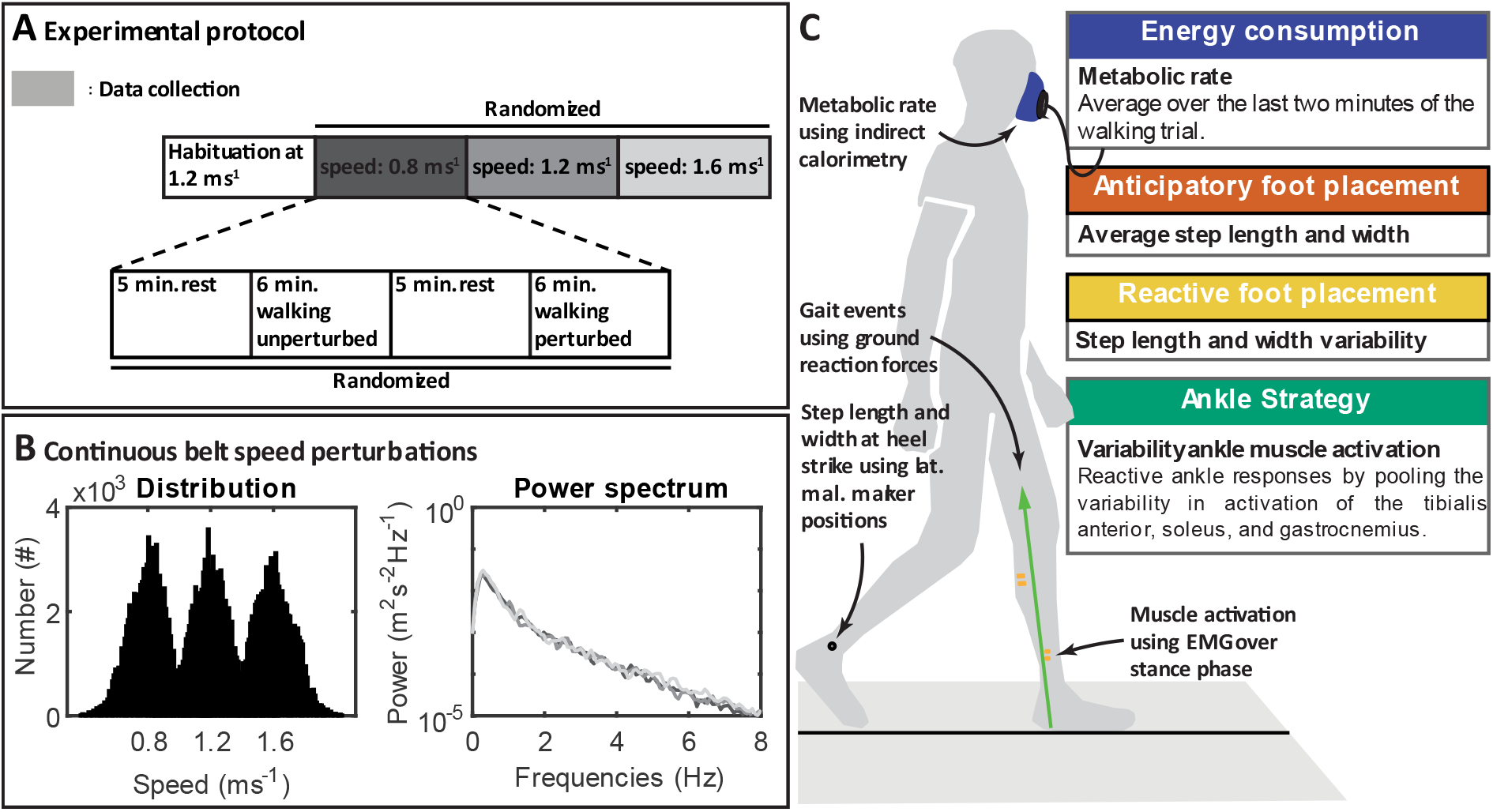
Summary of the experimental protocol. **A**. Schematic representation of order of experimental conditions and the randomization procedure. **B**. The histogram (left plot) depicts the number of discrete treadmill velocities (measured at 300Hz) within a certain velocity bin over the perturbation trials for each speed, while the power spectrum (right plot) shows the frequency content of the measured treadmill belt speed velocities for the three different speeds. **C**. Visualization of the relationship between experimental data and study outcomes (lat. mal. stands for lateral malleolus).

### Data acquisition

We measured oxygen consumption and carbon dioxide production to estimate metabolic energy consumption using the K5 metabolic measurement unit (COSMED, Rome, Italy). Marker and ground reaction force (GRF) data were acquired through the Vicon Giganet system (Oxford Metrics, Oxford, England) in combination with a Vicon camera system to measure the position of the lateral malleoli using retroreflective markers at 200 Hz to estimate step length and width at heel strike. Gait events were detected based on GRF, which were measured on the M-Gait instrumented treadmill at 1000 Hz. Ankle muscle activation patterns were measured using surface electromyography (EMG). EMG signals were sampled at 1000 Hz and measured using a bipolar electrode configuration (Cometa Wireless 32-channel EMG System, Cometa systems, Bareggio, Italy) or with the Delsys system (three last experiments - subjects 13, 21, and 22 – due to failure of the Cometa system; Trigno Wireless Biofeedback System with Avanti Sensors, Delsys, Natick, USA). EMG electrodes were placed over the tibialis anterior, gastrocnemius medialis, and soleus of the left leg after standard skin preparation (i.e. shaving and cleaning of the skin using alcohol).

### Perturbation protocol

During perturbation trials, balance was challenged by continuous random fore-aft treadmill belt speed variations. To construct the belt speed control signal for the perturbation trials, we adapted the procedure described in (15). The control signals for the velocity-controlled treadmill were generated in MATLAB and Simulink using software that was distributed with (15). First, we created pseudo-random belt acceleration using discrete Gaussian white noise with a standard deviation of 5 ms^−2^ and bounded between −15 and 15 ms^−2^. Then, we integrated the accelerations to generate a velocity perturbation signal. The velocity perturbation signal was high-pass filtered at 1.7 Hz to eliminate drift, bounded between 0 and 3.6 ms^−1^, and added to a constant speed of either 0.8, 1.2, or 1.6 ms^−1^ depending on the condition. Because we were interested in the effect of speed on balance control and balance-related energy consumption, we kept the perturbation magnitude constant over walking speeds. In contrast to (15), the velocity perturbation signal for all walking speeds was generated using the same inputs. We measured treadmill belt speed variation using the treadmill’s belt speed sensors. The standard deviation of measured velocities was approximately 0.13 ms^−1^. Conditions had different average belt speeds but similar frequency contents indicating a similar perturbation magnitude across walking speeds (Figure 1B).

### Data pre-processing

GRF marker data, and EMG signals were collected using Vicon’s Nexus. We labeled marker data in Nexus and processed data in MATLAB (R2023, MathWorks, Natick, USA). EMG data was band-pass filtered between 20 and 200 Hz, normalized to the median muscle activation over all unperturbed walking trials, and rectified. Rectified EMG, GRF, and marker data were low-pass filtered using a third order bidirectional Butterworth filter with a cut-off frequency of 13 Hz. Periods of missing marker data shorter than 0.1 seconds were linearly interpolated to account for marker occlusion.

In some EMG channels in subjects 3, 6, and 19 we registered large peaks, likely due to tension on the wires to the electrodes. We removed and linearly interpolated peaks with a prominence (amplitude from base to peak) larger than 20 standard deviations of the EMG over the entire trial by using MATLAB’s findpeaks algorithm. However, in some instances, removing peaks did not suffice to remove movement artifacts from the EMG data. Therefore, we excluded EMG data from these instances from further analysis. This was data in the perturbed trial from the gastrocnemius at speeds 1.2, and 1.6 ms^−1^ for subject 3, and from the tibialis anterior at speeds 0.8, 1.2, and 1.6 ms^−1^ for subject 6.

Heel strike and toe-off were defined as the moment at which the vertical component of the GRF under the respective foot increased above or decreased below 5% of body weight, respectively. We removed events at which both feet were in contact with the same force plate based on the vertical GRF under the foot. Gait events were included in further analysis if before heel strike and after toe-off 1) the vertical GRF was below 1% of the maximal GRF for 12% of the median stance duration, 2) the slope of the GRF after heel strike and before toe-off was sufficiently steep (i.e. average slope of the vertical GRF over 12% of median stance duration was at least 75% of the maximum force over the entire walking trial divided by the median stance phase duration), and 3) the vertical GRF reached at least half of the maximum value over the entire walking trial within the stance phase.

### Measurement equipment and outcomes

#### Metabolic energy consumption

We estimated metabolic rate from oxygen consumption and carbon dioxide production, measured by the K5, using the standard formula of Brockway (17). The energy consumption of the walking trials was represented by the average metabolic rate over the last two minutes of the trial. Subjects were asked not to consume caffeine (12 hours), nicotine (12 hours), or food (3 hours) before the experiment to prevent variability in measured signals unrelated to physical effort.

#### Step width and length

Step width and length were estimated by calculating the mediolateral and anteroposterior distance between markers on the lateral malleoli at heel strike of the leading leg. We used the difference in mean step width and length between perturbed and unperturbed walking at each speed to evaluate anticipatory changes to the gait pattern. We used the difference in standard deviation of step width and length between perturbed and unperturbed walking at each speed to evaluate changes in reactive foot placement.

#### Variability in ankle muscle activation

We expressed muscle activation (processed EMG signals) as a percentage of the single support phase of the left leg, i.e. the period from right toe-off to right heel strike when the right leg was in swing. First, we computed the standard deviation in muscle activation over steps at each percentage of the single support phase. Then, we computed the root mean square standard deviation over the single support phase for each muscle separately. Finally, we calculated the root mean square muscle activation standard deviations over ankle muscles to obtain one representative value for ankle muscle variability. We used the difference in ankle muscle variability between perturbed and unperturbed walking at each speed to evaluate changes in reactive ankle strategy.

We normalized metabolic rate to g^3/2^l^1/2^m, and step length and width to l to obtain all outcomes in dimensionless units (i.e. n.d.), where g is the gravity constant, l is leg length measured between the right lateral malleolus and posterior superior iliac spine, and m is the mass of the subject. Normalization factors for a subject with an average leg length and body weight were 2126 W for metabolic rate, and 0.93 m for step length and width.

### Statistical analysis

Statistical analysis was performed using R statistical software (R Foundation for Statistical Computing, Vienna, Austria). For testing our hypotheses, we used the difference in outcome variables between perturbed and unperturbed conditions as dependent variables in statistical models. We evaluated the effect of walking speed by fitting separate linear mixed-effect models on the change in metabolic rate, anticipatory foot placement (mean step width and step length), reactive foot placement (standard deviation in step width and length), and ankle strategy (ankle muscle activation variability) outcomes to test for the effect of walking speed with a fixed slope for speed and random intercept for each subject. In addition, we assessed the effect of perturbations on balance outcomes and metabolic rate for each walking speed separately using t-tests. When assumptions underlying parametric tests were violated, we used the Wilcoxon signed-rank test. We used R^2^ of the fixed and random effects, Cohen’s d for t-tests, and r scores for the Wilcoxon signed-rank test to quantify the effect sizes. We corrected for multiple comparisons to prevent Type-I error inflation due to the number of statistical tests that we performed using the Benjamini-Hochberg procedure. We considered probability values lower than 0.05 as statistically significant.

We explored the contribution of anticipatory, reactive foot placement, and ankle strategies to energy consumption using linear mixed-effect models by predicting between-subject variability in changes in metabolic rate due to perturbations from changes in balance outcomes due to perturbations. Here, we included speed as a categorical variable, balance outcomes as fixed effects, and a random offset per subject. Speed was included as a categorical variable as this reduced multicollinearity between variables compared to including speed as a numerical variable. We accounted for the interaction between speed and balance outcomes. Balance control outcomes that did not predict balance-related metabolic power were removed from the model through elimination based on the Akaike information criterion while retaining the random effect.

We tested whether age may have affected our results. As we aimed at collecting a dataset representative for the active population, the age range was large, i.e. 19 to 65 years old. Therefore, we explored whether adding age as a predictor in the linear mixed-effect models influenced the outcomes. Age did not affect the change in metabolic rate and balance control outcomes due to perturbations. However, we found some effects of age on the relationship between change in metabolic rate and changes in balance outcomes due to perturbations. Results from this analysis are described in the supplementary material (S2).

### Missing data

Missing ankle muscle variability data due to exclusion of some EMG data (see above) were replaced by substitute values for the statistical analysis by data imputation using the mice package in R (16). Substitute values were generated based on a two-level normal model with subject as a random and speed as a fixed effect. Standard deviations of activation for individual muscles were first imputed 30 times, generating multiple datasets. For each dataset, the average standard deviations over the three muscles were computed, and differences between perturbed and unperturbed walking were calculated to obtain values for ankle muscle variability for each imputed dataset. We then fitted linear mixed-effect models on each imputed data set and combined the results. In addition, we evaluated the effect of perturbations per walking speed using t-tests. While residuals of fitted linear-mixed effect models were normally distributed, the change in ankle muscle variability was not normally distributed for every walking speed. Instead of performing t-tests on the imputed dataset, we first bootstrapped 500 times and then imputed the bootstrapped datasets 30 times. We performed t-tests on each separate dataset and pooled the results across datasets.

### Departures from preregistration

Hypotheses and methods were preregistered before conducting this study (14). We decided to deviate from the preregistered document on different points. The most important departures from the pre-registration include the decision to apply data imputation for missing data, the use of pair-wise testing to assess the effect of perturbations for each speed independently, and to accommodate for multiple comparisons using the Benjamini-Hochberg procedure. A more elaborate description of the changes with respect to the pre-registered document can be found in the supplement (S1).

## Results

Statistical tests were performed after scaling outcomes to dimensionless units. For the sake of interpretability, we report in the text the corresponding values for the average subject, i.e. a subject with a leg length of 0.93 m and a body mass of 71.9 kg.

Subjects increased metabolic rate in perturbed compared to unperturbed walking trials and increases in metabolic rate were larger for slower than for faster walking speeds (Figure 2A). The metabolic rate of an average subject increased with 43 W at 0.8 ms^−1^, 30 W at 1.2 ms^−1^, and 21 W at 1.6 ms^−1^ (see Table 1), i.e. the metabolic rate associated with balance control decreased by 28 W for every 1 ms^−1^ increase in walking speed (slope=−0.013 m^−1^s, CI=−0.022 – −0.004 m^−1^s, p<0.001, R^2^=0.63).

**Table 1.** Difference in study outcomes between perturbed and unperturbed walking conditions at each walking speed. We described the central tendency, spread, and effect size using the mean, standard deviation (s.d.), cohens-d (d) for parametric tests and median, interquartile range (iqr), test statistic over the square root of the observations (r) for non-parametric tests. We scaled non-dimensional central tendencies (Non. dim. centr. ten.) to an average subject (Scaled centr. ten.). Probability values were corrected for multiple comparisons using the Benjamini-Hochberg procedure.

| Δmetabolic rate | Scaled centr. ten.<br>(median) | Non. dim. centr.<br>ten. (median) | Spread<br>(iqr) | P-value | Effect size<br>(r) |
| --- | --- | --- | --- | --- | --- |
| 0.8 ms<sup>-1</sup> | 43 W | 0.020 | 0.015 | <0.001 | 0.87 |
| 1.2 ms<sup>-1</sup> | 30 W | 0.014 | 0.013 | <0.001 | 0.83 |
| 1.6 ms<sup>-1</sup> | 21 W | 0.010 | 0.018 | 0.004 | 0.63 |
| Δmean step length | Scaled Centr. ten.<br>(median) | Non. Dim. Centr.<br>ten. (median) | Spread<br>(iqr) | P-value | Effect size<br>(r) |
| 0.8 ms<sup>-1</sup> | -0.034 m | -0.037 | 0.021 | <0.001 | 0.86 |
| 1.2 ms<sup>-1</sup> | -0.022 m | -0.023 | 0.029 | <0.001 | 0.76 |
| 1.6 ms<sup>-1</sup> | -0.025 m | -0.027 | 0.027 | <0.001 | 0.81 |
| Δmean step width | Scaled Centr. ten.<br>(mean) | Non. Dim. Centr.<br>ten. (mean) | Spread<br>(s.d.) | P-value | Effect size<br>(d) |
| 0.8 ms<sup>-1</sup> | 0.024 m | 0.026 | 0.017 | <0.001 | 1.55 |
| 1.2 ms<sup>-1</sup> | 0.018 m | 0.019 | 0.017 | <0.001 | 1.12 |
| 1.6 ms<sup>-1</sup> | 0.013 m | 0.014 | 0.011 | <0.001 | 1.19 |
| Δvar. step length | Scaled Centr. ten.<br>(median) | Non. Dim. Centr.<br>ten. (median) | Spread<br>(iqr) | P-value | Effect size<br>(r) |
| 0.8 ms<sup>-1</sup> | 0.031 m | 0.034 | 0.016 | <0.001 | 0.88 |
| 1.2 ms<sup>-1</sup> | 0.016 m | 0.017 | 0.007 | <0.001 | 0.88 |
| 1.6 ms<sup>-1</sup> | 0.008 m | 0.009 | 0.005 | <0.001 | 0.85 |
| Δvar. step width | Scaled Centr. ten.<br>(mean) | Non. Dim. Centr.<br>ten. (mean) | Spread<br>(s.d.) | P-value | Effect size<br>(d) |
| 0.8 ms<sup>-1</sup> | 0.003 m | 0.004 | 0.003 | <0.001 | 1.23 |
| 1.2 ms<sup>-1</sup> | 0.003 m | 0.003 | 0.003 | <0.001 | 0.99 |
| 1.6 ms<sup>-1</sup> | 0.003 m | 0.004 | 0.004 | <0.001 | 0.90 |

| Δvar. musc. act. | Non. Dim. Centr.<br>ten. (mean) | Non. Dim. Centr.<br>ten. (mean) | Spread<br>(s.d.) | P-value | Effect size<br>(d) |
| --- | --- | --- | --- | --- | --- |
| 0.8 ms<sup>-1</sup> | 1.12 | 1.12 | 0.72 | <0.001 | 1.55 |
| 1.2 ms<sup>-1</sup> | 0.81 | 0.81 | 0.50 | <0.001 | 1.63 |
| 1.6 ms<sup>-1</sup> | 0.51 | 0.51 | 0.35 | <0.001 | 1.44 |

**Figure 2.**
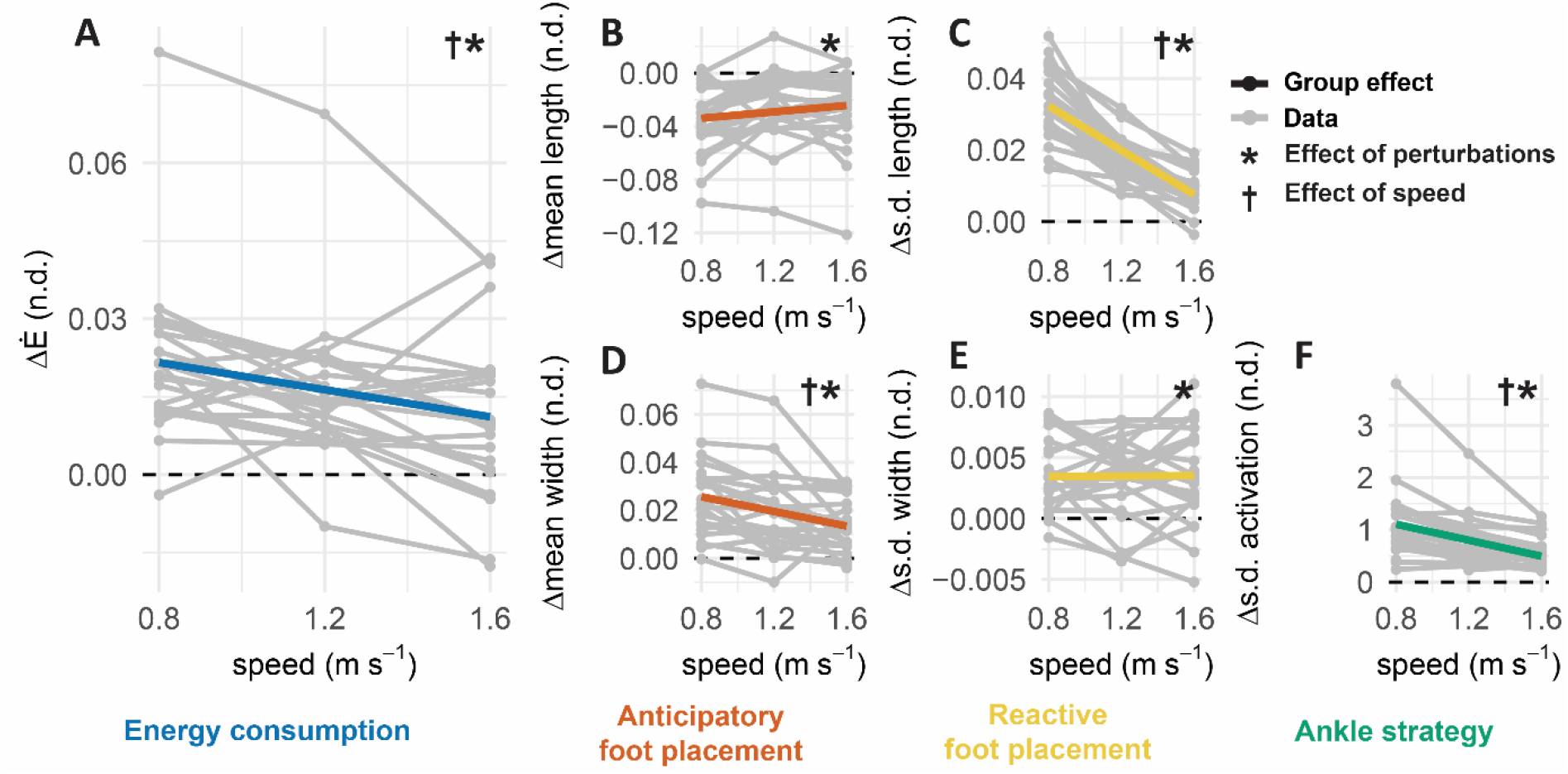
Effect of perturbations on metabolic rate and balance control outcomes as a function of walking speed. Graphs depict the difference between perturbed and unperturbed walking in study outcomes. **A**. Effect of perturbations on metabolic rate. **B**. Anticipatory adaptations in mean step length. **C**. Changes in the standard deviation (s.d.) of step length characterizing changes in reactive foot placement strategy. **D**. Anticipatory adaptations in mean step width. **E**. Changes in the standard deviation (s.d.) of step width characterizing changes in reactive foot placement strategy. **F**. Changes in ankle muscle activation variability characterizing changes in ankle strategy. All outcomes are non-dimensional (n.d.).

Subjects walked with shorter and wider steps in the perturbed compared to unperturbed condition, but step width increased less with increasing speed (see Figure 2B&D). Subjects decreased mean step length by 0.034 m at 0.8 ms^−1^, 0.022 m at 1.2 ms^−1^, and 0.025 m at 1.6 ms^−1^, and increased mean step width by 0.024 m at 0.8 ms^−1^, 0.018 m at 1.2 ms^−1^, and 0.013 m at 1.6 ms^−1^ in perturbed versus unperturbed walking (see Table 1). Adaptations in step width, but not step length, depended on walking speed. The average subject reduced adaptations in step width by 0.014 m^−1^s for every 1 ms^−1^ increase in walking speed (slope=−0.015 m^−1^s, CI=−0.023 – −0.008 m^−1^s, p<0.001, R^2^=0.62).

Step length and width were more variable in perturbed than in unperturbed walking but step length variability increased less with perturbations at faster speeds (see Figure 2C&E). For the average subject, step length variability increased in perturbed versus unperturbed walking by 0.031 m at 0.8 ms^−1^, 0.016 m at 1.2 ms^−1^, and 0.008 m at 1.6 ms^−1^ (see Table 1). Changes in step width variability were considerably smaller, the average subject increased step width variability by 0.003 m at 0.8, 1.2, and 1.6 ms^−1^ (see Table 1). Adaptations in step length variability decreased with walking speed by 0.029 m for every 1 ms^−1^ increase in speed for an average subject (slope = −0.031 m^−1^s, CI= −0.0352 – −0.0267 m^−1^s, p<0.001, R^2^=0.80).

Ankle muscle activation was more variable in perturbed than unperturbed walking but the increase in variability in response to perturbations was smaller at faster walking speeds. Variability in ankle muscle activation increased by 1.12 at 0.8 ms^−1^, 0.81 at 1.2 ms^−1^, and 0.51 at 1.6 ms^−1^ in perturbed compared to unperturbed walking (an increase of 1 corresponds to an increase equal to the median activity during unperturbed walking; see Table 1). Variability in ankle muscle activation increased less at fast than at slow walking speeds. Variability in ankle muscle activity decreased by 0.77 with every 1 ms^−1^ increase in speed (CI=−0.96 – −0.57 m^−1^s, p<0.001, R^2^=0.80).

Increases in energy consumption due to perturbations were variable across subjects, and this between-subject variability was partially explained by between-subject differences in changes in mean step length and the interaction with speed.Metabolic rate increased more if subjects reduced their step length more, and this was more pronounced at the slowest compared to the fastest walking speed. At 0.8 ms^−1^, metabolic rate increased by 8.1 W per cm decrease in mean step length for the average subject (or by 811 Wm^−1^, Table 2), while at 1.6 ms^−1^, metabolic rate increased by only 1.5 W per cm decrease in step length (or by 152 Wm^−1^). There were no differences between 0.8 and 1.2 ms^−1^, see Table 2.

**Table 2.** Changes in metabolic rate were partially explained by changes in mean step length. The linear mixed-effect model was fitted on dimensionless data. The model that predicted changes in metabolic rate after performing step reduction is described in the table below. Estimated effects can be found in the dimensionless effect column with corresponding confidence intervals (Non. dim. estimates). We scaled dimensionless estimates based on body weight and leg length of an average subject to aid the interpretation of the results (described in the Scaled estimates column).

| Predictors | Scaled estimates | Non. dim. estimates (CI) | p |
| --- | --- | --- | --- |
| Intercept | 20 W | 0.009 (0.0040 – 0.0147) | 0.001 |
| Δmean step length | -811 Wm<sup>-1</sup> | -0.353 (-0.4814 – -0.2247) | <0.001 |
| Δmean step length at 1.2 ms<sup>-1</sup> | 194 Wm<sup>-1</sup> | 0.084 (-0.0600 – 0.2286) | 0.247 |
| Δmean step length at 1.6 ms<sup>-1</sup> | 659 Wm<sup>-1</sup> | 0.287 (0.1519 – 0.4217) | <0.001 |
| Marginal R2 / Conditional R2 | 0.31 / 0.62 |  |  |

## Discussion

### In this study, we evaluated the metabolic cost of stabilizing walking in the sagittal plane across speeds and explored its relationship with balance control strategies

Random treadmill belt speed perturbations (standard deviation of velocity of 0.13 ms^−1^) elicited average increases in energy consumption of 16.7% at 0.8 ms^−1^, 9.8% at 1.2 ms^−1^, and 4.6% at 1.6ms^−1^, suggesting that there is a speed-dependent energetic cost associated with maintaining sagittal plane stability. Subjects adopted shorter and wider steps, and more variable foot placement and ankle muscle activation patterns during perturbed as compared to unperturbed walking, suggesting that subjects relied on both anticipatory adjustments of gait parameters and reactive foot placement and ankle strategies for balance control. Similar to changes in metabolic cost, changes in balance-related outcomes were smaller at faster than at slower speeds, suggesting that balance-related energetic costs are smaller when walking faster because walking faster is inherently more robust against disturbances. In addition, our exploratory analysis showed that between-subject variability in the increase in metabolic cost due to perturbations can – in part – be explained by between-subject variability in balance control strategies. Energy consumption increased less in perturbed versus unperturbed walking for subjects who adjusted their step length less, especially at slower speeds. Unperturbed walking requires balance control as well, and it is therefore likely that also during unperturbed walking, the energy cost of sagittal plane balance control is higher at slow than at faster speeds. This is especially relevant for individuals with mobility impairments. Many neuromusculoskeletal disorders lead to both a decrease in preferred walking speed (17–20) and alterations in balance control (21–24), which together might have an important contribution to the increased energetic cost observed in many neuromusculoskeletal disorders (18–20, 25).

### Although walking is thought to be inherently more robust against perturbations in the sagittal than in the frontal plane, we showed that there is also a considerable cost associated with sagittal plane balance control

Frontal plane walking balance is thought to require more active control – and thus energy – than sagittal plane walking balance. Therefore, most previous studies focused on the relationship between frontal plane balance control and energy consumption (2, 3) but O’Connor et al. (2) evaluated the effect of both mediolateral and fore-aft visual field perturbations. They found that mediolateral but not fore-aft visual perturbations increased the metabolic cost of walking. In contrast, we observed that the energetic cost of controlling sagittal plane walking balance can be considerable as well, i.e. energy consumption increased on average by 16.7% when walking at an average speed of 0.8 ms^−1^ with compared to without treadmill belt speed perturbations. It is likely that our perturbations challenged stability more than the visual perturbations applied by O’Connor et al. (2). Subjects might compensate for visual perturbations by relying more on other sensory inputs, whereas such a strategy is not possible for mechanical perturbations. In contrast to the belt-speed perturbations applied here, visual perturbations applied by O’Connor et al. (2) did not elicit alterations in foot placement. It is possible that participants relied on an ankle strategy when visually perturbed, but in that case, the lack of an increase in energy consumption would suggest that an ankle strategy is an efficient balance control strategy. A challenge with mechanical perturbations is that they might inject or dissipate mechanical energy and if so, this would complicate the interpretation of metabolic energy results. Here, we assume that no net work was performed by the perturbations on average because the speed was normally distributed around the mean speed.

### Subjects who relied more on anticipatory reductions of step length increased energy consumption more when perturbed

especially at slower speeds. In line with the relationship between increases in metabolic cost and anticipatory adaptations in the walking pattern observed here, Ahuja and Franz (2022) found that both metabolic cost and step width decreased during adaptation to visual perturbations in the frontal plane (3). Whereas studies investigating the energy cost associated with anticipatory balance control are scarce, the energy cost of voluntary alterations in step length and width has been extensively studied (26, 27). When people are asked to reduce step length and thus increase step frequency while maintaining the same walking speed, they consume more energy (28). The observation that energy consumption increases when subjects walk at higher or lower frequencies than their preferred frequency has contributed to the hypothesis that humans select gait patterns that minimize energy consumption (29). It is unclear whether humans select a balance strategy that minimizes energy consumption or prioritize robustness against perturbations over energetic cost. In other words, anticipatory reductions in step length might reduce the need for reactive balance strategies, which also come at an energetic cost, and it is possible that humans select a combination of anticipatory and reactive balance strategies by optimizing the overall metabolic cost. Between-subject variability in this trade-off might then be explained by differences in the neuro-musculoskeletal system. Alternatively, anticipatory adjustments might increase robustness to perturbations, and some individuals might prefer this increased robustness even if it comes at an increased metabolic cost.

### It remains challenging to determine contributions of different balance control strategies to energy consumption based on the experiment performed here

Our results suggest that anticipatory adaptations to the gait pattern contributed most to the energetic cost of stabilizing walking. Yet, individuals who rely more on anticipatory strategies may rely less on reactive strategies. This potential interdependence between anticipatory and reactive balance control hinders the attribution of changes in energy consumption for balance control to either an increased reliance on anticipatory or a decreased reliance on reactive balance strategies. We used a principal component analysis to explore the interdependence of our five balance control measures (mean step length and width, variability of step length, step width and muscle activity; Supplementary Figure S1). All five principal components contributed considerably to explaining between-subject differences in changes in balance outcomes (at least 10% at 0.8 m/s, 4% at 1.2 m/s, and 7% at 1.6 m/s), indicating that no outcomes were redundant. However, at the slowest walking speed, the component explaining most of the variability (31%) couples larger changes in anticipatory step length adjustments to smaller changes in reactive step length variability and ankle muscle variability. This suggests that subjects who relied more on anticipatory balance control strategies relied less on reactive balance responses. This coupling might have prevented us from detecting relationships between the reliance on reactive balance responses and changes in metabolic energy consumption. To further explore relationships between metabolic cost and balance control strategies, a larger sample (since outcomes are independent to some degree) or experiments that manipulate the participants’ ability to use specific balance control strategies are required.

### The metabolic cost of controlling balance is higher at slower than at faster speeds suggesting that more active balance control is required at slow speeds

At faster speeds, both changes in balance outcomes and increases in metabolic cost were smaller than at lower speeds. This suggests that increasing speed improves the robustness against perturbation. Hak and colleagues (5) also concluded that walking at faster speeds is inherently more robust against perturbations by evaluating the margin of stability across a combination of speeds and step frequencies in unperturbed walking. Faster speed and higher step frequencies increased the stability margin in the backward direction, while higher step frequency also improved the mediolateral stability margin. As far as we know, this study is the first to show the importance of walking speed when studying the contribution of balance control to energy consumption.

### The higher energetic cost of balance control at slow versus fast walking speeds has important implications for many mobility-impaired individuals who walk slower

Mobility impairments, e.g. due to aging or lower-limb amputation, are often associated with lower walking speeds (17, 19, 21). Our results suggest that the slower walking speed might amplify age- or disease-related balance control problems and increase metabolic cost. Here, we used perturbations to assess the energetic cost of walking but balance control is also required during unperturbed walking. Sensorimotor processes that underlie human motor control are thought to be stochastic (30), meaning that stability during walking is continuously challenged by noisy sensory information and motor controls. In addition, the environment is uncertain as well, e.g. the floor is never exactly level and this is especially true when walking outside. Our findings might, therefore generalize to unperturbed walking. Indeed, when walking is externally stabilized using springs, metabolic cost and variability in foot placement decrease (1). When balance control is compromised by age or disease (31–33), balance control strategies and, thereby also the energetic cost of balance control might change. This might explain why we found interactions with age when exploring the relationship between increases in metabolic cost and balance strategies (Supplementary Table S1). We found an interaction effect of step width variability and age on increases in metabolic energy. A larger increase in step width variability was related to lower balance-related energy consumption but more so in young than in older adults. In a prior study, we observed that older adults relied more on a reactive stepping strategy and modulated ankle muscle activity less than young adults when perturbed (11). An increase in step width variability in older adults might thus indicate a shift from reactive ankle to stepping strategy that comes with a metabolic penalty. The observed interaction with age might indicate that there are different interactions between reactive step width adjustments and other balance strategies in young and older adults. It is possible that young adults who rely more on a reactive stepping strategy rely less on anticipatory adjustments and that this comes with a metabolic advantage. Similarly, individuals with neuro-musculoskeletal disorders might use different balance strategies, affecting their metabolic cost. For example, individuals with a transtibial amputation can no longer use an ankle strategy on the amputated side and might have to rely more on anticipatory and reactive stepping strategy. This might contribute to the increased metabolic cost that is often observed in individuals with a transtibial amputation.

In conclusion, continuous treadmill speed perturbations increased energy consumption associated with balance control more at slow than at fast walking speeds and this might be explained by walking being more robust at faster than at slower walking speeds. Our results suggest that the use of anticipatory balance control strategies contributed most to the energetic cost of walking balance control, especially at slow walking speeds. These findings are especially relevant for explaining the increased metabolic cost of individuals with mobility impairments, who often walk slower and have altered balance control.

## Supplementary materials

### S1. Deviations from methods described in the preregistered document

Deviations from the methods described in the preregistered document were small (14).

1. We decided to use the marker positions of the lateral malleoli to estimate step length and width instead of using the OpenSim pipeline. Using the Opensim pipeline would have been more time-consuming with little gain in accuracy.
2. In the preregistered document, we described that missing data would be excluded from further analysis. In the final approach, we decided to use data imputation to handle missing data. As a result, we excluded less data from the analysis. For example, our measure for ankle muscle variability consisted of pooled standard deviations over three muscles. When data from one of the muscles was missing, data was replaced by imputation and combined with the available data from the other muscles to obtain the outcome for ankle muscle activation variability instead of excluding all data from further analysis.
3. We normalized the metabolic rate and stepping outcomes to the dimensions of the individuals before performing statistical tests. By accounting for between-subject variability in body dimensions, we reduced the variability in balance outcomes (e.g. step length and changes in step length are expected to increase with leg length) that were not related to between-subject differences in balance control strategies.
4. In addition to linear mixed-effect models, we decided to test the effect of perturbations per walking speed separately using t-tests and Wilcoxon signed-rank tests. These tests are straightforward to perform and interpret compared to deriving the effect of perturbations per speed from linear mixed-effect models. The decision to test the effect of perturbation for each speed increased the number of comparisons. To accommodate for the number of comparisons, we performed a Benjamini-Hochberg correction on test outcomes.
5. We used a cut-off frequency of 13 instead of 20 for the low-pass filter on rectified EMG, marker, and GRF data.

### S2. Interaction between balance outcomes and age

We included individuals from a large age range, i.e. between 19 and 65 years old. We explored whether age influenced the outcomes by adding age as a predictor in the linear mixed-effect models. We did not detect an effect of age on metabolic rate, anticipatory foot placement, reactive foot placement, and ankle strategy outcomes but we observed an interaction effect of age on the relationship between between-subject variability in increases in metabolic rate and changes balance control outcomes upon perturbations.

Our analysis accounting for age confirmed the results described in the main manuscript. Metabolic rate increased more if subjects reduced their step length more and this was more pronounced at the slowest compared to the fastest walking speed. At 0.8 ms^−1^, metabolic rate increased by 7.0 W per cm decrease in step length for the average subject (or by 702 Wm^−1^, Table 2), while at 1.6 ms^−1^, metabolic rate increased by only 0.6 W per cm decrease in step length (or by 63 Wm^−1^). There were no differences between 0.8 and 1.2 ms^−1^, see Table 2. In addition, we found that metabolic rate increased less if subjects increased step width variability more and that this effect was smaller as age increased. Metabolic rate decreased by 21 W per cm increase in step width variability (or by 2197 Wm^−1^, Table S1). This decrease in metabolic rate with more variable step width was smaller when subjects were older by 194 Wyear^−1^m^−1^. While we found no relationship between step length variability and metabolic rate, we did find an interaction with age. Metabolic rate decreased with increased step length variability depending on age by 24 Wyear^−1^m^−1^.

**Table S1.**
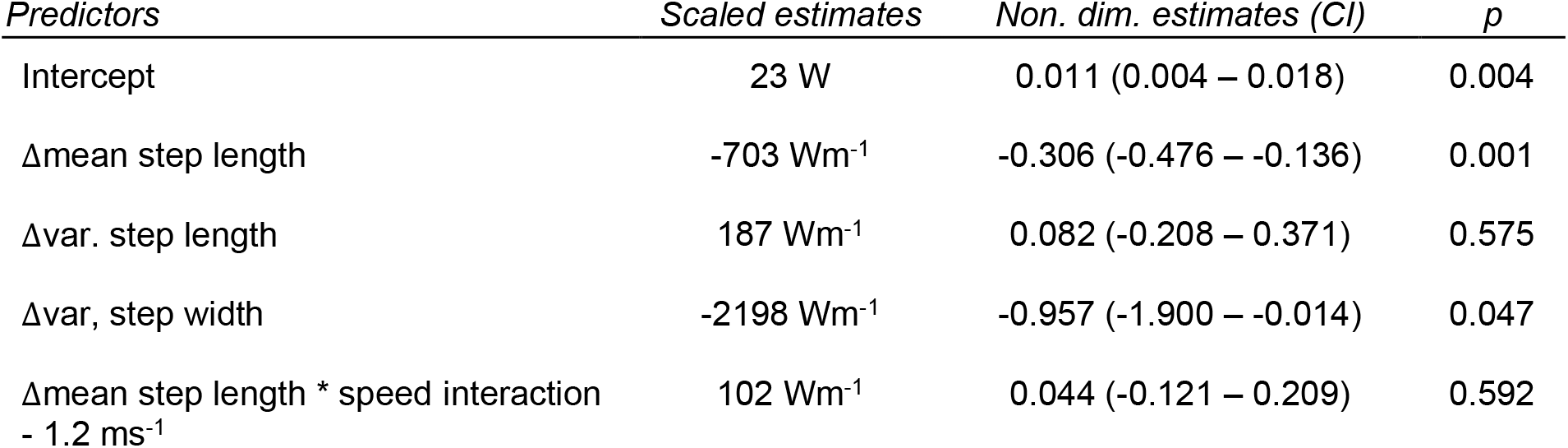
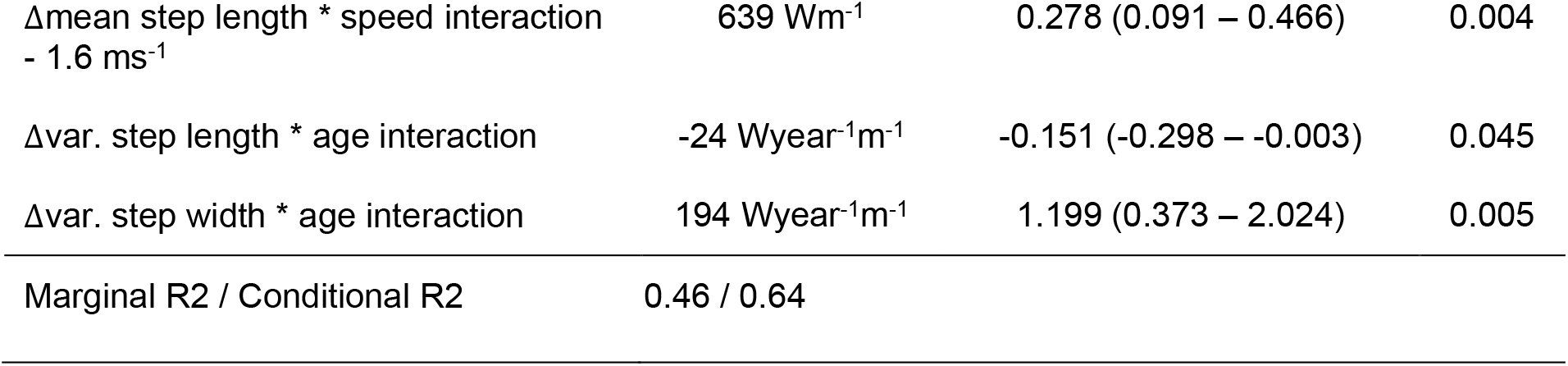
Changes in metabolic rate were partially explained by adaptations in a subset of balance control outcomes when we included the effect of age. The linear mixed-effect model was fitted on dimensionless data. The model that predicted changes in metabolic rate best after performing step reduction, is described in the table. Estimated effects can be found in the dimensionless effect column with corresponding confidence intervals (Non. dim. estimates). We scaled dimensionless estimates to a subject of average body weight, leg length, and age to aid the interpretation of the results (described in the Scaled estimates column). We mean-centered age before including it as a variable in the linear mixed-effect model, the interaction with age describes the effect of age with respect to the mean age of 39-years-old.

| Predictors | Scaled estimates | Non. dim. estimates (CI) | p |
| --- | --- | --- | --- |
| Intercept | 23 W | 0.011 (0.004 – 0.018) | 0.004 |
| Δmean step length | $-703 \text{ Wm}^{-1}$ | $-0.306 (-0.476 - -0.136)$ | 0.001 |
| Δvar. step length | $187 \text{ Wm}^{-1}$ | $0.082 (-0.208 - 0.371)$ | 0.575 |
| Δvar, step width | $-2198 \text{ Wm}^{-1}$ | $-0.957 (-1.900 - -0.014)$ | 0.047 |
| Δmean step length * speed interaction<br>- $1.2 \text{ ms}^{-1}$ | $102 \text{ Wm}^{-1}$ | $0.044 (-0.121 - 0.209)$ | 0.592 |
| Δmean step length * speed interaction<br>- 1.6 ms<sup>-1</sup> | 639 Wm<sup>-1</sup> | 0.278 (0.091 – 0.466) | 0.004 |
| Δvar. step length * age interaction | -24 Wyear<sup>-1</sup>m<sup>-1</sup> | -0.151 (-0.298 – -0.003) | 0.045 |
| Δvar. step width * age interaction | 194 Wyear<sup>-1</sup>m<sup>-1</sup> | 1.199 (0.373 – 2.024) | 0.005 |
| Marginal R2 / Conditional R2 | 0.46 / 0.64 |  |  |

### S3. Principal component analysis of interdependence between balance outcomes

We explored whether we indeed needed all five balance outcomes to describe between-subject variability in balance control strategies by performing a principal component analysis. We performed the principal component analysis on changes in balance outcomes in perturbed versus unperturbed walking at each speed separately. All five components contributed considerably to the between-subject variability indicating that we cannot reduce the set of balance outcomes. Principal components were to some extent similar across speeds. Notably, the component explaining the largest amount of the variability at the two slowest speed (31% and 38%) and the second largest amount of variability at the fastest speed (29%) (Component 1, Figure S1) has large positive loadings on mean step length and large negative loadings on step length variability. This indicates that a large portion of inter-subject variability can be explained by a different trade-off between anticipatory and reactive adjustments in step length. At the slowest speeds, the component with large positive loadings on ankle muscle activation variability and large negative loadings on step length variability (Component 3, see Figure S1) explained 16% and 20% of the variability. This suggests that another important part of the between-subject variability can be explained by different trade-offs between reactive ankle and stepping strategies. The size of the loadings on ankle muscle activation variability decreases with increasing speed, suggesting that a reduced reliance on the ankle strategy at higher speeds as previously observed (4) is consistent across subjects.

**Figure S1.**
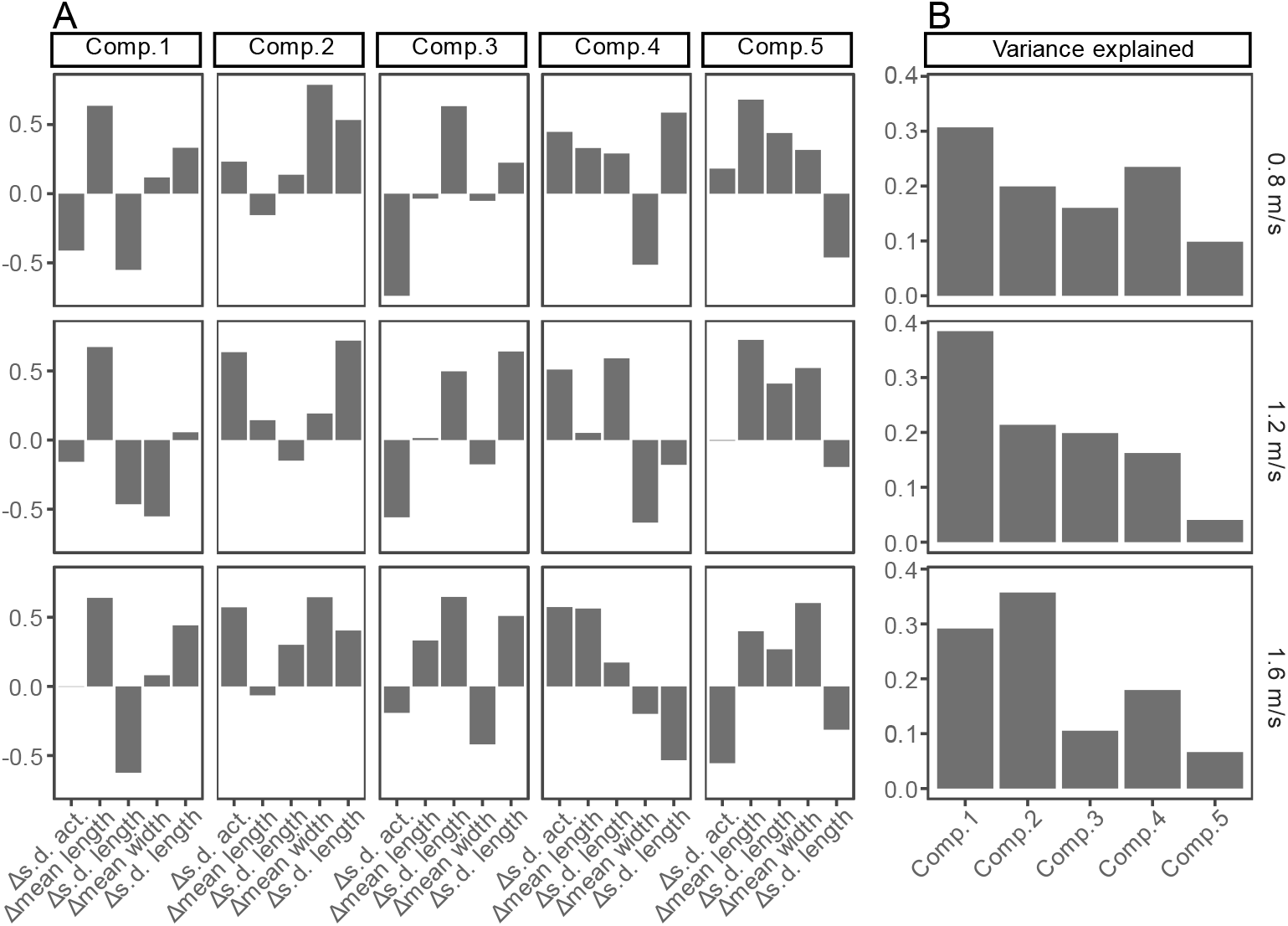
Principal component analysis of between-subject variability at different speeds. **A**. The loadings for each of the five components, in the columns, at different walking speeds, in the rows, with s.d. act. for the standard deviation of ankle muscle activation, s.d. length for the standard deviation of step length, and s.d. width for the standard deviation of step width. **B**. The variance explained by the components at each speed. Components were sorted to match the component vectors of the 1.2 ms^−1^ walking speed by comparing vectors using the inner product. Acknowledgments

We would like to thank all participants as well as the colleagues who dedicated their time to being pilot subjects. We are grateful for the support that Ruben Hillewaere provided over the course of the experiments.

## Grants

The project was funded by a FWO fellowship (1SF7322N) received by WM.

## Disclosures

The authors declare that the research was conducted in the absence of any commercial or financial relationships that could be construed as a potential conflict of interest.

## Author contributions

FDG, RR, MA, and WM developed the study concept and designed the experiments. After the design of the study, MA and WM preregistered the study. WM acquired and analyzed the data. FDG and WM wrote the manuscript. FDG, RR, MA, and WM interpreted the results and edited the manuscript. All authors contributed to the article and approved the submitted version.

## Notes

### Competing Interest Statement

The authors have declared no competing interest.

## References

1. Donelan JM, Shipman DW, Kram R, Kuo AD. Mechanical and metabolic requirements for active lateral stabilization in human walking. Journal of Biomechanics 37: 827–835, 2004. doi: 10.1016/j.jbiomech.2003.06.002.

2. O’C Xu HZ, Kuo AD. Energetic cost of walking with increased step variability. Gait Posture 36: 102–107, 2012. doi: 10.1016/j.gaitpost.2012.01.014.

3. Ahuja S, Franz JR. The metabolic cost of walking balance control and adaptation in young adults. Gait & Posture 96: 190–194, 2022. doi: 10.1016/j.gaitpost.2022.05.031.

4. Afschrift M, Groote FD, Jonkers I. Similar sensorimotor transformations control balance during standing and walking. PLOS Computational Biology 17: e1008369, 2021. doi: 10.1371/journal.pcbi.1008369.

5. Hak L, Houdijk H, Beek PJ, Van Dieë JH. Steps to take to enhance gait stability: The effect of stride frequency, stride length, and walking speed on local dynamic stability and margins of stability. PLoS ONE 8, 2013. doi: 10.1371/journal.pone.0082842.

6. Espy DD, Yang F, Bhatt T, Pai Y-C. Independent influence of gait speed and step length on stability and fall risk. Gait & Posture 32: 378–382, 2010. doi: 10.1016/j.gaitpost.2010.06.013.

7. Voloshina AS, Kuo AD, Daley MA, Ferris DP. Biomechanics and energetics of walking on uneven terrain. Journal of Experimental Biology 216: 3963–3970, 2013. doi: 10.1242/jeb.081711.

8. O’C Kuo AD. Direction-Dependent Control of Balance During Walking and Standing. Journal of Neurophysiology 102: 1411–1419, 2009. doi: 10.1152/jn.00131.2009.

9. Swart SB, den Otter R, Lamoth CJC. Anticipatory control of human gait following simulated slip exposure. Sci Rep 10: 9599, 2020. doi: 10.1038/s41598-020-66305-1.

10. Yang F, Kim J, Munoz J. Adaptive gait responses to awareness of an impending slip during treadmill walking. Gait & Posture 50: 175–179, 2016. doi: 10.1016/j.gaitpost.2016.09.005.

11. Afschrift M, van Deursen R, De Groote F, Jonkers I. Increased use of stepping strategy in response to medio-lateral perturbations in the elderly relates to altered reactive tibialis anterior activity. Gait & Posture 68: 575–582, 2019. doi: 10.1016/j.gaitpost.2019.01.010.

12. Vlutters M, van Asseldonk EHF, van der Kooij H. Center of mass velocity-based predictions in balance recovery following pelvis perturbations during human walking. Journal of Experimental Biology 219: 1514–1523, 2016. doi: 10.1242/jeb.129338.

13. Vlutters M, van Asseldonk EHF, van der Kooij H. Reduced center of pressure modulation elicits foot placement adjustments, but no additional trunk motion during anteroposterior-perturbed walking. Journal of Biomechanics 68: 93–98, 2018. doi: 10.1016/j.jbiomech.2017.12.021.

14. Muijres W, Afschrift M, Ronsse R, De Groote F. What is the contribution of balance control to the metabolic cost of walking with sagittal plane treadmill speed perturbations? OSF n.d., 2024. doi: 10.17605/osf.io/t6zjq.

15. Moore JK, Hnat SK, van den Bogert AJ. An elaborate data set on human gait and the effect of mechanical perturbations. PeerJ 2015, 2015. doi: 10.7717/peerj.918.

16. Buuren S, Groothuis-Oudshoorn C. MICE: Multivariate Imputation by Chained Equations in R. Journal of Statistical Software 45, 2011. doi: 10.18637/jss.v045.i03.

17. Hermodsson Y, Ekdahl C, Persson BM, Roxendal G. Gait in male trans-tibial amputees: A comparative study with healthy subjects in relation to walking speed. Prosthet Orthot Int 18: 68–77, 1994. doi: 10.3109/03093649409164387.

18. Theunissen K, Plasqui G, Boonen A, Brauwers B, Timmermans A, Meyns P, Meijer K, Feys P. The Relationship Between Walking Speed and the Energetic Cost of Walking in Persons With Multiple Sclerosis and Healthy Controls: A Systematic Review. Neurorehabilitation and Neural Repair 35: 486, 2021. doi: 10.1177/15459683211005028.

19. Schrack JA, Simonsick EM, Chaves PHM, Ferrucci L. The Role of Energetic Cost in the Age-Related Slowing of Gait Speed. Journal of the American Geriatrics Society 60: 1811–1816, 2012. doi: 10.1111/j.1532-5415.2012.04153.x.

20. Brehm M-A, Nollet F, Harlaar J. Energy Demands of Walking in Persons With Postpoliomyelitis Syndrome: Relationship With Muscle Strength and Reproducibility. Archives of Physical Medicine and Rehabilitation 87: 136–140, 2006. doi: 10.1016/j.apmr.2005.08.123.

21. Hak L, Dieën JH van, Wurff P van der, Prins MR, Mert A, Beek PJ, Houdijk H. Walking in an Unstable Environment: Strategies Used by Transtibial Amputees to Prevent Falling During Gait. Archives of Physical Medicine and Rehabilitation 94: 2186–2193, 2013. doi: 10.1016/j.apmr.2013.07.020.

22. Gates DH, Scott SJ, Wilken JM, Dingwell JB. Frontal plane dynamic margins of stability in individuals with and without transtibial amputation walking on a loose rock surface. Gait & Posture 38: 570–575, 2013. doi: 10.1016/j.gaitpost.2013.01.024.

23. Mohamed Suhaimy MSB, Okubo Y, Hoang PD, Lord SR. Reactive Balance Adaptability and Retention in People With Multiple Sclerosis: A Systematic Review and Meta-Analysis. Neurorehabil Neural Repair 34: 675–685, 2020. doi: 10.1177/1545968320929681.

24. Ofran Y, Schwartz I, Shabat S, Seyres M, Karniel N, Portnoy S. Falls in Post-Polio Patients: Prevalence and Risk Factors. Biology (Basel) 10: 1110, 2021. doi: 10.3390/biology10111110.

25. Ettema S, Kal E, Houdijk H. General estimates of the energy cost of walking in people with different levels and causes of lower-limb amputation: a systematic review and meta-analysis. Prosthetics and Orthotics International 45: 417–427, 2021. doi: 10.1097/PXR.0000000000000035.

26. Maxwell Donelan J, Kram R, Arthur D. K. Mechanical and metabolic determinants of the preferred step width in human walking. Proceedings of the Royal Society of London Series B: Biological Sciences 268: 1985–1992, 2001. doi: 10.1098/rspb.2001.1761.

27. Kuo AD, Donelan JM, Ruina A. Energetic Consequences of Walking Like an Inverted Pendulum: Step-to-Step Transitions. Exercise and Sport Sciences Reviews 33: 88, 2005.

28. Holt KG, Hamill J, Andres RO. Predicting the minimal energy costs of human walking. Medicine & Science in Sports & Exercise 23: 491, 1991.

29. Alexander RM. Optimization and gaits in the locomotion of vertebrates. Physiological Reviews 69: 1199–1227, 1989. doi: 10.1152/physrev.1989.69.4.1199.

30. van Beers RJ, Baraduc P, Wolpert DM. Role of uncertainty in sensorimotor control. Philos Trans R Soc Lond B Biol Sci 357: 1137–1145, 2002. doi: 10.1098/rstb.2002.1101.

31. Ivers RQ, Cumming RG, Mitchell P, Attebo K. Visual impairment and falls in older adults: the Blue Mountains Eye Study. Journal of the American Geriatrics Society 46: 58– 64, 1998. doi: 10.1111/J.1532-5415.1998.TB01014.X.

32. Lord SR, Menz HB. Visual Contributions to Postural Stability in Older Adults. Gerontology 46: 306–310, 2000. doi: 10.1159/000022182.

33. Pijnappels M, van der Burg JCE, Reeves ND, van Dieën JH. Identification of elderly fallers by muscle strength measures. European Journal of Applied Physiology 102: 585– 592, 2008. doi: 10.1007/s00421-007-0613-6.

